# The atypical filament assembly underpins the inflammasome-independent functions of IFI16

**DOI:** 10.1101/2025.01.02.631089

**Authors:** Archit Garg, Ewa Niedzialkowska, Jeffrey J. Zhou, Jasper Moh, Edward H. Egelman, Jungsan Sohn

## Abstract

Inflammasomes trigger cell death upon sensing various intracellular maladies. These supra-structures relay upstream signals by sequentially assembling architecturally congruent filaments via their pyrin domains (PYDs). Interferon Inducible Protein 16 (IFI16) is an innate immune sensor that detects dysregulated nucleic acids. Once presumed as an inflammasome receptor due to its PYD, the role of IFI16 has been much more appreciated in other innate immune pathways such as regulating interferon production and viral replication restriction. Here, a cryo-EM structure of the filament assembled by the PYD of IFI16^(PYD)^ shows a helical architecture distinct from inflammasome PYD filaments. *In silico Rosetta* interaction energy calculations suggest that the helical architecture of the IFI16^PYD^ filament is incompatible with those assembled by the central inflammasome adaptor ASC and its interacting partners. Cellular experiments further support that IFI16 fails to interact with ASC. Together, we provide the structural basis for the inflammasome-independent functions of IFI16.

## Introduction

Inflammasomes are supramolecular signaling platforms that play vital roles in host innate defense against various intracellular maladies, which include organelle damage, tumor formation, and pathogen invasion ^1^. A unique aspect of inflammasomes is that signal transduction is executed by sequentially assembling filamentous oligomers ^2–4^. That is, upon detecting molecular signatures arising from intracellular maladies (e.g., viral DNA and dysfunctional organelles), the inflammasome receptors undergo oligomerization and assemble filaments using their pyrin-domains (PYDs) ^1–5^. These filaments then provide a template for the polymerization of the PYD of the central inflammasome adaptor termed ASC, (ASC^PYD^, apoptosis associated speck forming protein containing caspase-recruiting domain (CARD)) ^2–5^. ASC is a bipartite protein composed of a PYD and a CARD, and the ASC^PYD^ filament assembly results in its CARD filament formation, which then leads to the polymerization and activation of caspase-1 ^1,4^. The activated protease then executes key host defense processes such as maturation of pro-inflammatory cytokines and pyroptotic cell death ^6,7^.

Both PYD and CARD belong to the death-domain (DD) superfamily, which are about 100 amino-acids (a.a.) long, 6-helix bundles ^8^. Multiple cryo-EM studies revealed that all known PYD filaments share the same helical architecture (hexameric base, six protomers at the axial pole) ^3–5,9–11^, while the CARD-subfamily filaments display their own common architecture (tetrameric base, four protomers at the axial pole) ^12–18^. Given that several upstream PYDs (and CARDs) can signal through one common downstream filament ^9,11,18,19^, such subfamily-specific congruent architectures provide an efficient framework for signal integration and transduction ^19^. Moreover, recent studies have identified that self-assembly of DD filaments occur bidirectionally, while the recognition between two signaling partners occur unidirectionally ^3,5^. Nonetheless, it is still unclear how these signaling filaments recognize and distinguish their interaction partners. It also remains unknown whether there exist PYD- and/or CARD filament assemblies whose structures deviate from their subfamily specific architectures, what such noncanonical assemblies would entail in dictating their signaling partners.

Interferon inducible protein 16 (IFI16) is a versatile innate immune sensor that detects dysregulated nucleic acids ^20^, with its most prominent function being assembling filaments on unchromatinized double-stranded (ds)DNA ^20–24^. Upon assembly, IFI16 partakes in a wide range of innate immune responses such as accentuating cGAS-STING-dependent interferon signaling pathways, viral replication restriction, and even DNA damage responses by regulating chromatin dynamics ^25–30^. Nevertheless, how and by what signaling partners IFI16 interacts with in its various innate immune functions remain largely speculative. For example, a long-standing conundrum about IFI16 is whether it directly activates ASC-dependent inflammasomes. That is, IFI16 belongs to the family of Absent-in-Melanoma-2 (AIM2)-like receptors (ALRs), which consist of a PYD and one or two DNA-binding HIN200 domains (hematopoietic interferon inducible nuclear antigen with 200 a.a.) ^1,31^. It has been proposed that AIM2 is the only ALR that directly activates inflammasomes upon detecting cytosolic dsDNA by inducing the filament assembly of ASC ^31–33^. However, albeit lacking evidence for direct interactions, there have been a few cellular studies suggesting that IFI16 induces ASC-dependent inflammasome formation in specific viral infection cases involving Kaposi sarcoma herpes virus (KSHV) and HIV ^34,35^. Of note, we recently found that full-length IFI16^(FL)^ fails to induce the polymerization of ASC^PYD^ regardless of bound nucleic acids using recombinant proteins (e.g., dsDNA, dsRNA, ssDNA, and ssRNA) ^24^, supporting the view that IFI16 does not directly interact with ASC^PYD^. Nonetheless, the structural mechanisms that allow IFI16 to operate outside of ASC remain unknown.

We present here a cryo-EM structure of the IFI16^PYD^ filament at 3.3 Å, which reveals that it assembles into a helical architecture distinct from all other known PYD filaments ^3–5,10^. This atypical assembly is largely accomplished by the unique amino acids (a.a.) compositions in two areas that alter the secondary structures of the IFI16^PYD^ monomer. *Rosetta* interface energy calculations suggest that the helical architecture of the IFI16^PYD^ filament is indeed incompatible with ASC^PYD^ and its upstream receptors. Cellular experiments also support our structure and *in silico* predictions that IFI16 does not directly interact with ASC. Together, our results provide the structural basis not only for the versatile functions of IFI16 outside of inflammasomes, but also for understanding how filamentous signaling platforms regulate their specificities by altering helical architectures.

## Results

### The cryo-EM structure of the IFI16^PYD^ filament

Although essential for understanding its diverse biological function ^25–30^, the structure of the IFI16^PYD^ filament remains unknown. After extensive optimization, we identified conditions that allow recombinant IFI16^PYD^ to auto-assemble into filaments (Figure 1A). The average power-spectrum from cryo-EM images of 256-pixel long nonoverlapping filament segments showed that the IFI16^PYD^ filament is a three-start helix with 134.6° rotation and 5.6 Å rise, displaying no rotational symmetry (Figure 1B). Because the structure of IFI16^PYD^ monomer is unknown, we used AlphaFold ^36^ for model building. The resolution of the final model was 3.3 Å according to the gold standard method (Figure 1C-D). The diameter of the outer rim is 85 Å and the inner cavity is at 22 Å (vs. 94 Å and 25 Å in ASC^PYD ref. 4^) (Figure 1E). The assembly of the IFI16^PYD^ filament is also mediated by the six interfaces that are commonly found in PYDs and CARDs (Types 1-3 ref. ^4^; Figure 1F). The Type 1a:1b interface is mediated by side-chain interactions including polar and hydrophobic residues (Figure 1F-b), and Type 2a:2b and Type 3a:3b interfaces include mostly polar and ionic side-chain interactions (Figure 1F-c/d). We also noted side-chains we previously identified important for filament assembly and dsDNA-mediated oligomerization of full-length IFI16 (e.g., Leu^28^ and Ile^46^) ^23^.

**Figure 1.**
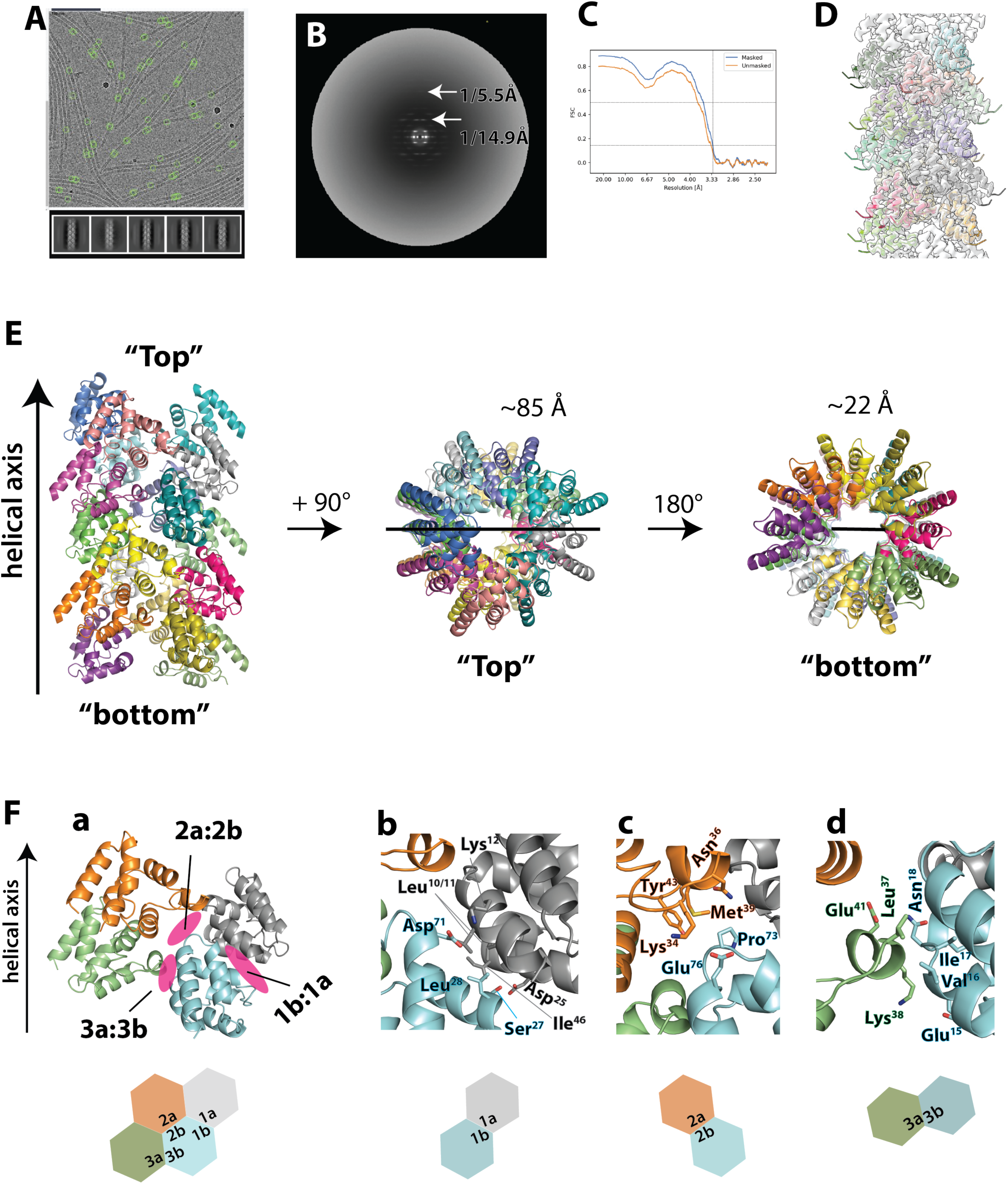
The cryo-EM structure of the IFI16^PYD^ filament. **(A)** Cryo-electron micrographs of the IFI16^PYD^ filaments. 2D classes are shown below. **(B)** The average power spectrum of IFI16^PYD^ filaments. **(C)** The FSC curve of the cryo-EM map of the IFI16^PYD^. The dotted lines indicate the 0.143 threshold for the resolution. **(D)** The IFI16^PYD^ filament model built into the cryo-EM map. **(E)** The atomic model of the IFI16^PYD^ filament. **(F)** The three unique filament interfaces are shown along with the cartoon representation below. **(b-d)** side chains that mediate protein-protein interactions at each unique interface are shown.

Unlike the inflammasome filaments assembled by ASC^PYD^ and its interacting partners such as AIM2^PYD^, our structure shows that the IFI16^PYD^ filament lacks the C3 symmetry and displays different helical parameters (Figure 2A) ^3–5,10^. Moreover, instead of the six protomers that constitute the filament base of ASC^PYD ref. 4^, the base of the IFI16^PYD^ filament was composed of five protomers (i.e., hexagon vs. skewed pentagon; Figure 2A-B). As a consequence, the diameters of both outer and inner rims of the IFI16^PYD^ filament were smaller than those of the ASC^PYD^ filament (Figure 2B). Additionally, the top and bottom surfaces of both IFI16^PYD^ and ASC^PYD^ filaments were consistently electro-negative and positive, respectively; however, the charges were concentrated around the outer rim in the IFI16^PYD^ filament, while it was either evenly distributed (negative) or following the C3 symmetry (positive) for the ASC^PYD^ filament, resulting a mismatch (Figure 2C).

**Figure 2.**
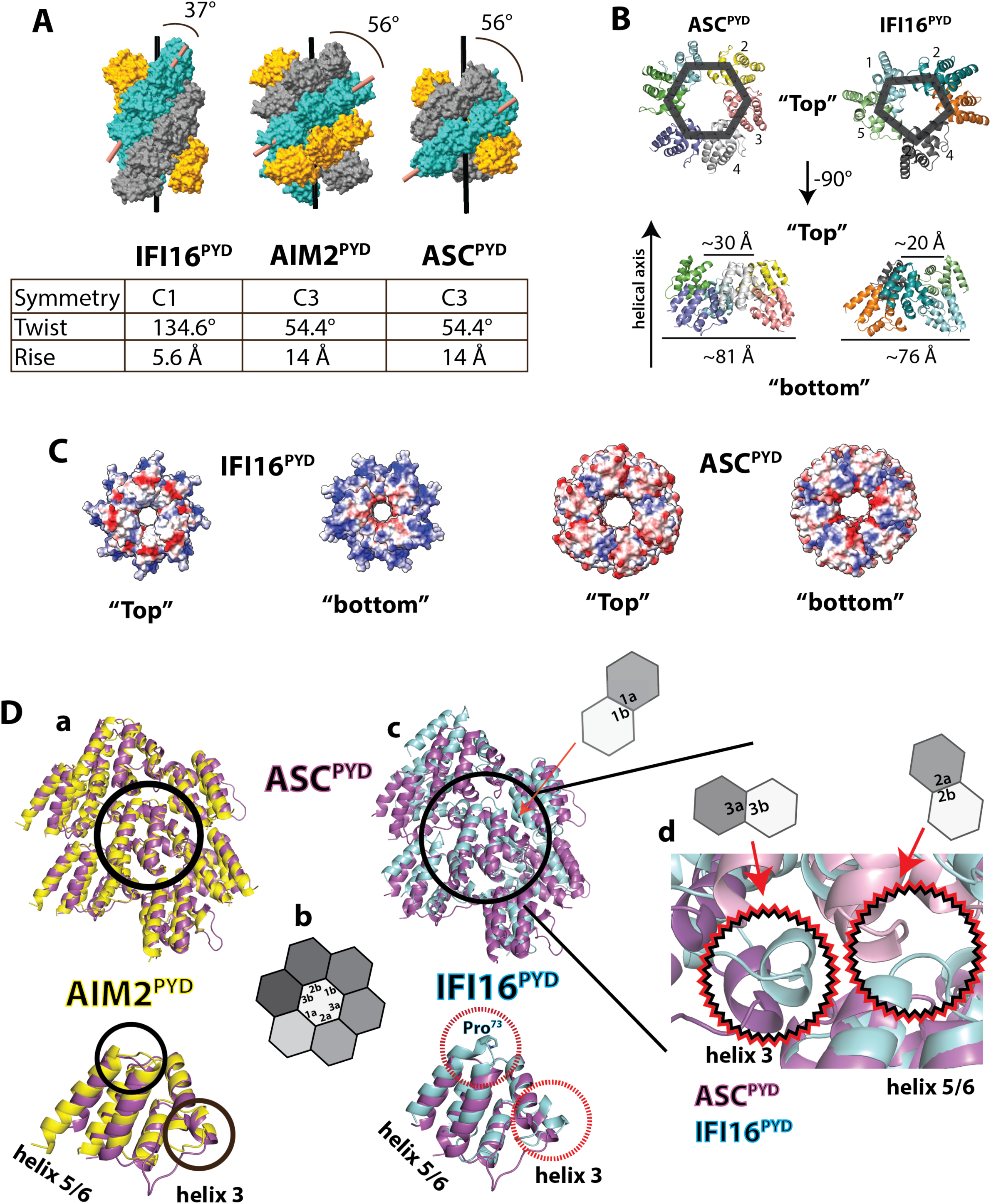
The architecture of the IFI16^PYD^ filament is not compatible with the ASC^PYD^ filament. **(A)** Comparing the IFI16^PYD^ filament to those assembled by AIM2^PYD^ (PDB ID: 7k3r) and ASC^PYD^ (PDB ID: 3j63). The angle between the central axis to the repeating helical strand is indicated. Key helical parameters are listed in the table. **(B)** Top- and side views comparing the ASC^PYD^ and IFI16^PYD^ filaments. A hexagon and a pentagon are shown to indicate the number of subunits at the filament base. **(C)** The electrostatic distribution of ASC^PYD^ and IFI16^PYD^ filaments. **(D) (a)** An overlay between AIM2^PYD^ and ASC ^PYD^ filament “honeycombs” and monomers. The middle subunit is indicated with a black circle in the filament. The regions that AIM2^PYD^ and ASC^PYD^ do not deviate in the monomers are also indicated with black circles. **(b)** A schematic showing the “honeycomb” representation of PYD filaments. Each unique surface is noted in the center subunit and the surrounding subunits are colored in different shades of gray. **(c)** An overlay of IFI16^PYD^ and ASC ^PYD^ filament honeycombs and monomers. The middle subunit is indicated with a black circle in the honeycomb. The regions that deviate in the two proteins are indicated with dotted red circles in the monomers. **(d)** A zoom-in view of the two key areas that would clash between IFI16^PYD^ and ASC^PYD^ filament assemblies. Hexagons in **(c)** and **d)** show the location of clash/deviations using the interface identifiers shown in **(b)**.

To further compare the architecture of IFI16^PYD^ filament to the ASC^PYD^ filament, we then generated the “honeycomb” side-views of PYD filaments in which the middle subunit makes all repeated contacts with surrounding subunits for assembly (Figure 2D; Figure 2D-b shows the corresponding honeycomb diagram with the six interfaces labeled; see also ref. ^3,37^). As expected, ASC^PYD^ and AIM2^PYD^ filaments aligned well with the monomers showing minimal deviations (RMSD: 0.8 Å; Figure 2D-a). By contrast, although the IFI16^PYD^ monomer was highly homologous to ASC^PYD^ (RMSD: 1.5 Å), the filament architecture deviated drastically despite aligned at the center (Figure 2D-c).

Further examination indicated two key structural differences in the IFI16^PYD^ monomer that would clash with ASC^PYD^-like assembly. For instance, compared to ASC^PYD^, the short helix 3 that mediates the Type 3 interface protrudes from the center of the protein by ∼ 6 Å in IFI16^PYD^ (dotted red circle in Figure 2D-c). Additionally, the loop between helices 5 and 6 in the Type 2 interface also extends from the core by ∼ 5 Å due to Pro^73^ that shortens the helix 5 and lengthens the helix 6 (dotted red circle in Figure 2D-c). These differences in IFI16^PYD^ would then clash with the ASC^PYD^-like assembly in the Type 3 and Type 2 interfaces (Figure 2D-d), also causing the Type 1 interface to misalign (Figure 2D-c, red arrow).

To examine whether such differences are induced by filament assembly or the a.a. composition intrinsic to IFI16^PYD^, we then compared it to the PYD of MNDA (88% sequence identity to IFI16; Supplementary Figure 1A-a). We noted that MNDA^PYD ref. 38^ closely resembles IFI16^PYD^ (RMSD: 0.7 Å; Supplementary Figure 1A-b). The a.a. sequence of the helix 3 and connecting loops are poorly conserved in ALRs and ASC (Supplementary Figure 1A-a), yet the helix 3 conformations of IFI16 and MNDA were still similar (Supplementary Figure 1A-b). These observations suggest that the unique helical architecture of the IFI16^PYD^ filament is largely built into its primary a.a. sequence (filament assembly likely stabilizes the flexible helix 3).

### Rosetta *in silico* analyses suggest that IFI16 is incompatible with inflammasome PYD-like assembly and *vice versa*

Our structure of the IFI16^PYD^ filament provides a compelling explanation as to why it would not directly interact with ASC. We then used *Rosetta* energy calculations to further examine the compatibility between IFI16^PYD^ and other filaments involved in the inflammasome pathway ^3,37^. Here, for self-assembly, the IFI16^PYD^ honeycomb displayed favorable energy scores seen from other inflammasome PYD filaments ^3,37^ (Figure 3A-a/b). These scores were also markedly more favorable compared to when we modeled IFI16^PYD^ into the AIM2^PYD^ honeycomb in our previous study (Figure 3A-b vs. Supplementary Figure 1B; see also ref. ^37^). Conversely, when we modeled ASC^PYD^ and AIM2^PYD^ using the IFI16^PYD^ honeycomb as a template, the resulting scores were dramatically less favorable compared to the scores from their native structures (Figure 3B and Supplementary Figure 1C; see also ref ^3,37^). Next, to compare the interface energetics between ASC^PYD^ and either ALR, we replaced the center subunit of the receptor filament honeycombs with ASC^PYD^ (Figure 3C). Here, the interface energy scores arising from ASC^PYD^ in the IFI16^PYD^ shell were markedly less favorable compared to the scores from AIM2^PYD^•ASC ^PYD^ (Figure 3C-a/b). Consistent with our *in silico* analysis, when co-transfected into HEK293T cells as eGFP/mCherry-tagged proteins, the AIM2^PYD^ filament co-localized with the ASC^PYD^ filament, while the IFI16^PYD^ filament failed to do so (Figure 3D). These results consistently suggest that the atypical helical architecture of the IFI16^PYD^ filament precludes its interaction with ASC^PYD^.

**Figure 3.**
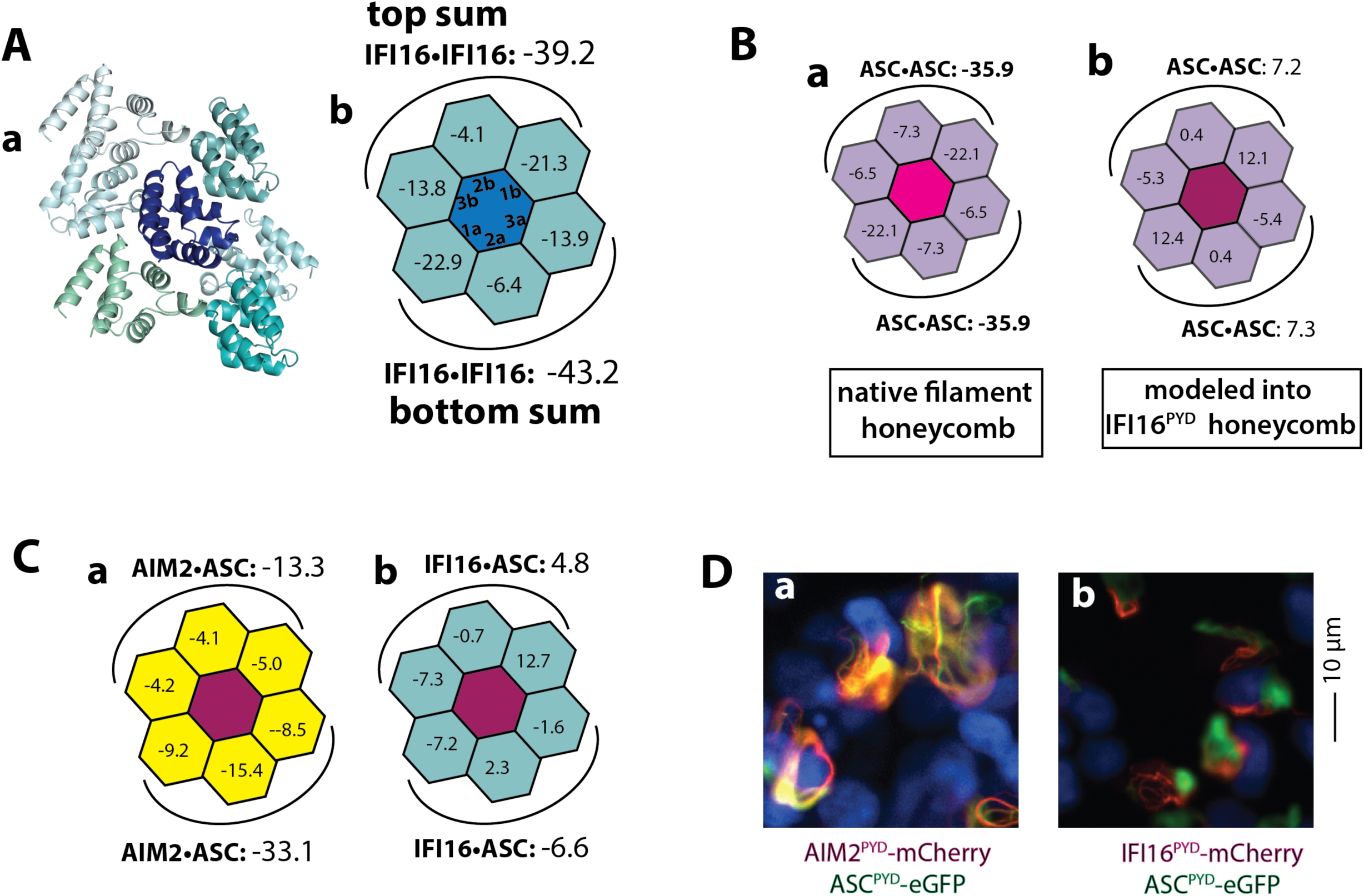
IFI16^PYD^ does not directly interact with ASC^PYD^. **(A) (a)** The honeycomb sideview of the IFI16^PYD^ filament. The center protomer (dark blue) makes all six unique contacts with surrounding protomers for assembly. **(b)** The corresponding honeycomb diagram of IFI16^PYD^ filament with *Rosetta* interface energy units (*reus*). The sum of reus from the top and bottom halves are also shown. **(B)** The *reus* of the ASC^PYD^ honeycombs modeled when using **(a)** the native filament (PDB: 3j63) vs. **(b)** that of IFI16^PYD^. **(C)** The resulting *reus* when ASC^PYD^ was modeled into either the (**a**) AIM2^PYD^ (PDB ID: 7k3r) or **(b)** IFI16^PYD^ honeycomb. **(D)** Fluorescence microscope images of HEK293T cells co-transfected with eGFP-tagged ASC^PYD^ and either mCherry-tagged AIM2^PYD^ or IFI16^PYD^. Blue: DAPI.

Some helical filaments can adapt different symmetries during assembly. For example, bacterial Pili filaments can assume the C5 symmetry or assemble without any rotational symmetry ^39^. Of note, it was previously reported that the AIM2^PYD^ filament can assemble with no rotational symmetry, albeit this architecture was caused by the N-terminal GFP-tag interfering with the native C3 symmetry ^40^. Like the IFI16^PYD^ filament, five subunits constitute the base of the GFP-AIM2^PYD^ filament, but not six seen from the untagged filament ^3,40^. We thus compared the GFP-AIM2^PYD^ structure to the IFI16^PYD^ filament. Here, when we overlayed the two honeycombs, it was apparent that the overall assembly of the IFI16^PYD^ filament is still different (Supplementary Figure 1D-a). Moreover, the energy scores resulting from modeling IFI16^PYD^ using the GFP-AIM2^PYD^ honeycomb were much less favorable than its native assembly (Figure 3A-b vs. Supplementary Figure 1D-b; −40 vs. −13 for half-sum scores; see also ref. ^37^). Our observations again suggest that the architecture of the IFI16^PYD^ filament is unlikely to be accessible by inflammasome PYDs such as AIM2.

## Discussion

Inflammasomes play vital roles in host innate defense against various maladies ranging from pathogen invasion to the exposure to genotoxic stresses ^1^. A key aspect of inflammasome pathways is the signal transduction by sequentially assembling homologues filaments ^1–5^. For instance, there are more than a dozen upstream receptors that recognize different types of molecular signatures associated with intracellular calamities (e.g., cytosolic dsDNA, malformed trans-Golgi network, and viral RNA) ^1^. These initially divergent pathways then converge at the assembly of the ASC filament ^1^. This is possible because, regardless of sensing mechanisms, all inflammasome receptors contain PYDs, and despite their a.a. sequence divergence in PYDs, it had been postulated that they all assemble into architecturally congruent helical filaments to the ASC^PYD^ filament ^1–5^. We and others found that within these homologous supra-structures there exist side-chain interactions that promote bidirectional self-assembly and the unidirectional recognition between upstream receptor PYD filaments and ASC^PYD ref. 3,5^. Here, we determined the cryo-EM structure of the IFI16^PYD^ filament, which revealed its unique helical assembly distinct from all other known PYD filaments. Our structure provides mechanistic basis for its versatile innate immune functions outside of ASC-dependent inflammasomes ^20,21,25–29^.

There are four well-known ALRs: AIM2, IFI16, IFIx, and MNDA ^20,31^. The primary sequence of AIM2 (both PYD and HIN200) is most divergent from the other three ALRs (i.e., Supplementary Figure 1A), and IFI16 has an extra dsDNA-binding HIN domain ^20,31^. It is well-established that the primary function of AIM2 is to assemble the ASC-dependent inflammasome ^31,33^, while the other three ALRs represent a rare case for PYD-containing proteins whose biological functions are much more appreciated outside of inflammasomes. Most notably, IFI16 accentuates the cGAS-STING pathway, regulates viral latency and replication, suppresses oncogene expression, and controls chromatin dynamics upon DNA double-strand break ^20,21,25–29,34^. Despite such versatile roles, how exactly IFI16 recognizes its signaling partners, and whether it can directly induce ASC-dependent inflammasome assembly remain controversial ^31–35^. Combined with our previous observation in which recombinant IFI16^FL^ fail to regulate the polymerization of ASC^PYD^ regardless of bound nucleic acids ^24^, our results here further indicate that the IFI16 filament by itself does not directly participate in inflammasome assembly. Instead, we envision two possibilities for IFI16 to still induce inflammasome formation via ASC. First, it is possible that there exist yet unidentified viral- or endogenous proteins that can bridge IFI16 and ASC. Alternatively, post-translational modification of IFI16 triggered by specific infectious conditions can alter the helical architecture of IFI16 filament, which might allow it to interact with ASC. Future studies directing these possibilities will illuminate the role of IFI16 in ASC-dependent inflammasome activation.

PYDs belong to the DD superfamily and often found in proteins involved in inflammatory signaling pathways ^8^. There are two well-known subclasses of PYDs. First, the vast majority of PYD-containing proteins are thought to act as the upstream receptor for ASC-dependent inflammasome formation ^1^. The other class consists of pyrin-only-proteins (POPs) that interfere and regulate the assembly of various inflammasome PYD filaments ^37,41^. The structure of the IFI16^PYD^ filament presented here marks the third subclass of the PYD family where it assembles into architecturally distinct filaments for their non-inflammasome functions. Indeed, there is a subset of NLRPs whose role is to downregulate the activation of ASC-dependent inflammasomes ^42,43^. Determining the structures of these PYD filaments will further delineate how the architectural diversity underpins the cellular functions of PYD filaments.

## Materials and Methods

### Cloning, expression and purification of recombinant IFI16^PYD^

The human IFI16^PYD^ construct (residues 1-94) was cloned into a pET21b vector with a C-terminal His_6_-tag and transformed into *Escherichia coli* BL21^DE3^ cells. Protein expression was induced around OD_600_: 0.5 using 0.2 mM isopropyl β-D-1-thiogalactopyranoside at 18 °C for 16-18 h. The cells were resuspended homogenously and lysed by sonication in lysis buffer containing 25 mM NaPO_4_, pH 7.8, 400 mM NaCl, 8 mM β-mercaptoethanol, 40 mM imidazole and 10% glycerol. Cell lysate after centrifugation was loaded on a pre-equilibrated Ni^+2^-NTA column and further purified using Superdex 75 16/600 size-exclusion column (Cytiva). Fractions containing IFI16^PYD^ were pooled, concentrated to 95 μM and stored in 20 mM HEPES pH 7.4, 400 mM NaCl, 1 mM EDTA, 1 mM DTT and 5% glycerol at −80 °C.

### Cryo-EM sample preparation and data collection

The IFI16^PYD^ filament sample was buffer exchanged to 40 mM HEPES pH 7.4, 160 mM NaCl, 1 mM EDTA and 1 mM DTT and concentrated to 73 μM. 5 μl sample was applied to glow discharged Lacey grids followed by auto-blotting for 4 s at 100% humidity and room temperature, which were then plunge frozen in liquid ethane using FEI Vitrobot Mark IV. Frozen grids were clipped using the clipping station and cryo-EM data were collected at the Beckman Center for cryo-EM (Johns Hopkins School of Medicine) using the Thermo Scientific Glacios TEM operating at 200 kV and equipped with the Falcon 4i direct electron detector. 3,140 movies with a total dose of ∼40 electrons/Å^2^ were collected at a pixel size of 1.165 Å/pixel and a defocus range of −0.7 to −3.0 μm.

### Helical reconstruction and model building

The data was processed using CryoSPARC v4 ^44^. The 3,140 movies, with a sampling of 1.165 Å/px, were subjected to “patch motion correction”. The defocus values and astigmatism of the micrographs were determined by CTF estimation (patch CTF estimation) for the aligned full dose micrographs. A total of 1,072 micrographs were selected based on relative ice thickness and CTF fit resolution for subsequent image processing. Filament tracer was used for segment picking from long filaments. The CTF-corrected micrographs were used for the segment extraction, with 256 px long boxes, with a shift of 40 Å between adjacent boxes. These particles were subjected to 2D classification to identify classes containing single filaments and to remove bad segments. An averaged power spectrum was generated from rotationally aligned segments and used for determining the helical symmetry parameters. The selected particles were then processed with the helical refinement for the final reconstruction after the helical parameters (an azimuthal rotation of 134.6° and an axial rise of 5.6 Å per subunit) converged. The resolution of the final reconstruction was estimated by the FSC between two independent half maps, which showed 3.3 Å at FSC = 0.143.

We used AlphaFold ^36^ to generate a model based on closely related MNDA^PYD^ structure (PDB ID:5H7Q ^38^) and docked into the IFI16^PYD^ cryo-EM map by rigid body fitting, and then manually edited the model in Chimera ^45^and Coot ^46^. The IFI16^PYD^ filament model was validated with MolProbity ^47^.

### Cell culture and imaging

For cell imaging, each construct was cloned into the pCMV6 vector containing a C-terminal mCherry or eGFP tag. Indicated plasmids (600 or 1200 ng) were (co)-transfected into HEK293T cells at ∼70% confluency using lipofectamine 3000 (Invitrogen). After 18 hours, cells were washed with 1x PBS, fixed with 4% paraformaldehyde followed by staining with NucBlue reagent (Invitrogen) and mounted on glass slides using ProLong Gold antifade reagent (Invitrogen). Images were captured using the Cytation 5 plate reader equipped with a fluorescent microscope then processed with the Gen5 software (Agilent).

### Rosetta simulation

The InterfaceAnalyzer script in Rosetta was used to determine the interaction energy at individual interfaces of the honeycomb as described in ^ref. 3,37^. We used the cryo-EM structures of IFI16^PYD^ (this study), ASC^PYD^ (PDB: 3j63), AIM2^PYD^ (PDB: 7k3r), GFP-AIM2^PYD^ (PDB: 6mb2) filaments to generate corresponding honeycombs.

## Acknowledgement

We thank the Sohn lab members for discussion. We also thank Drs. Seamus Morrone and Mariusz Matyzewski for their prior efforts. This work was supported by NIH 35GM145363, NSF MCB1845003, and Johns Hopkins Accelerator Awards to J.S; NIH 35GM122510 to E.H.E.

---

**Supplementary Figure 1.**
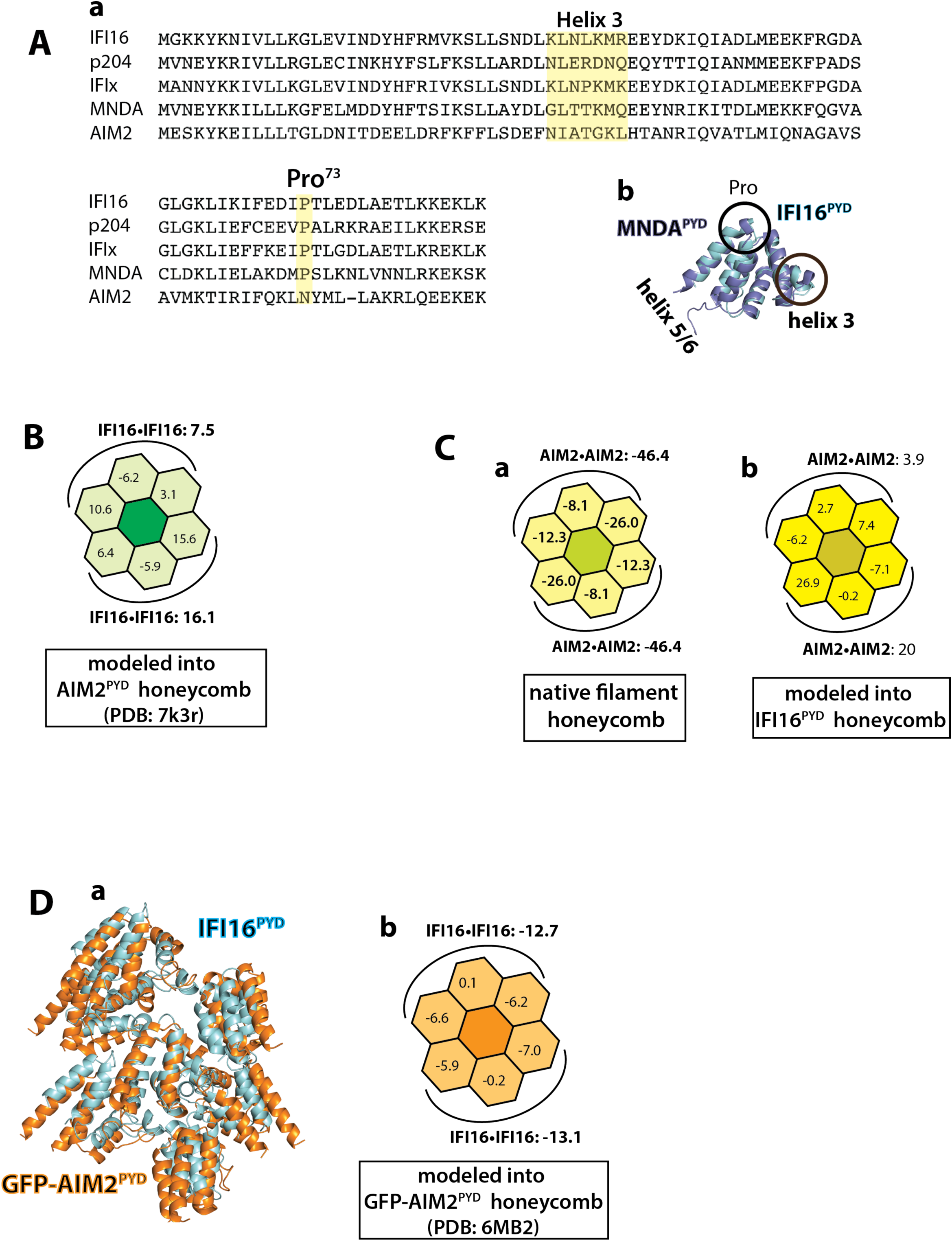
**(A)** (**a**)The sequence alignment of ALRs. (p204: mouse IFI16). Helix 3 and Pro73 (IFI16) are highlighted. (**b**) An overlay between MNDA^PYD^ (PDB ID: 5h7q) and IFI16^PYD^ monomers. **(B)** The *reus* when the IFI16^PYD^ filament was modeled using the AIM2^PYD^ honeycomb. **(C)** The *reus* of the AIM2^PYD^ honeycombs when modeled using **(a)** the native filament (PDB: 7k3r) vs. **(b)** that of IFI16^PYD^. **(D) (a)** An overlay between GFP-AIM2^PYD^ (PDB ID: 6mb2) and IFI16 ^PYD^ filament “honeycombs”. **(b)** The *reus* when the IFI16^PYD^ filament was modeled using the GFP-AIM2^PYD^ honeycomb.

